# Decreased food intake contributes to elevated insulin-responsiveness in pre-clinical cancer cachexia

**DOI:** 10.64898/2026.03.12.711318

**Authors:** Emma Frank, Kaspar W. Persson, Zakarias Kefil Jensen Ogueboule, Tang Cam Phung Pham, Jonas R. Knudsen, Lykke Sylow, Steffen H. Raun

## Abstract

**Purpose:** Cancer cachexia is a life-threatening complication of advanced malignancies, driven by anorexia and profound systemic metabolic reprogramming. Insulin action in skeletal muscle is markedly impaired in patients with cancer and may contribute directly to cachexia pathogenesis. However, the interplay between reduced nutrient intake and cancer-associated metabolic rewiring in cachexia remains poorly defined. Clarifying this relationship is essential for identifying the fundamental drivers of cachexia and for developing effective therapeutic strategies.

**Methods:** We assessed metabolic rewiring by glucose tolerance test and isotopic tracers to determine muscle insulin-stimulated glucose uptake in male cachectic and non-cachectic C26- and KPC-tumor-bearing, as well as mice towards C26 cachectic mice.

**Results:** Cachectic C26-tumor-bearing mice displayed reduced body weight, lean, and fat mass, and food intake (-20%, -15%, -75%, -40%, respectively). Cachectic C26- and KPC-tumor mice showed improved glucose tolerance compared to non-cachectic mice, correlating inversely with tumor size. Ex vivo insulin-stimulated glucose uptake was elevated in soleus (+78%) and extensor digitorum longus (+35%) muscle from cachectic C26-cancer mice compared to non-cachectic and control mice. This increase was associated with enhanced AKT signaling. This was phenocopied in pair-fed non-tumor-bearing mice to match the food intake of cachectic mice, where glucose tolerance, insulin-stimulated glucose uptake ex vivo, and AKT signaling were all enhanced by food restriction.

**Conclusions:** Our findings suggest that enhanced skeletal muscle insulin responsiveness in cachectic tumor-bearing mice is due to anorexia-induced adaptations, highlighting AKT signaling as a key node connecting nutrient status to muscle glucose metabolism in cancer cachexia.

**Highlights:**

- C26 and KPC cancer-induced weight loss (cachexia) increases glucose tolerance in mice
- Insulin responsiveness is increased in cachectic, but not in non-cachectic, tumor–bearing mice.
- Lowered food intake drives elevated muscle insulin responsiveness in cachectic mice

## 1. Introduction

Cancer cachexia is a multifactorial metabolic syndrome characterized by involuntary loss of skeletal muscle mass, with or without concomitant fat loss, due to reduced appetite and systemic metabolic rewiring. Affecting up to 80% of patients with advanced cancer, cachexia contributes substantially to morbidity, treatment intolerance, mortality, and impairs quality of life[1,2]. Systemic metabolic rewiring in cancer involves reduced insulin responsiveness in skeletal muscle[3,4], possibly contributing to the pathogenesis of cachexia. However, the interplay between reduced food intake and cachexia-associated metabolic reprogramming remains poorly defined. Yet, it is central in delineating the mechanisms that drive cachexia and in identifying treatment targets.

Skeletal muscle, being a major site of glucose disposal under insulin-stimulated conditions, is likely to play a central role in the metabolic disturbances associated with cancer. Clinically, cancer is often associated with systemic insulin resistance and impaired glucose tolerance in patients[5–7] to a degree that is similar to the insulin resistance observed in patients with type 2 diabetes[8]. Accordingly, patients with cancer have a higher risk of developing type 2 diabetes after their diagnosis[9,10]. In the preclinical setting, weight-stable LLC-tumor-bearing mice display impaired insulin tolerance and skeletal muscle insulin resistance[11–13]. Because insulin is an anabolic hormone, insulin resistance might contribute to cancer cachexia. This has been indicated by some preclinical evidence showing that insulin resistance precedes cachexia and that cachexia can be delayed by insulin sensitizers in a rat model of cachexia[14,15]. Moreover, patients with cancer and diabetes display a greater weight loss compared to patients with cancer but without diabetes[16]. Yet, cachectic patients did not exhibit insulin resistance, in contrast to the general cancer patient population[7] and non-cachectic preclinical mouse tumor-bearing mice[11,12,17], as indicated by a recent systematic review and meta-analysis [18]. Moreover, at basal conditions, cachectic mouse muscle was recently shown to have glucose hypermetabolism[19]. Thus, it remains unclear whether cachexia is associated with altered insulin responsiveness, and potential mechanisms linking cancer cachexia with insulin action are unresolved.

A key contributor to cachexia that could influence insulin action is the reduced food intake associated with cancer cachexia. Anorexia in patients and tumor-bearing mice exacerbates weight loss by limiting energy and nutrient availability, accelerating muscle and fat wasting, but it is not the sole driver of cachexia[20–22]. However, reduced caloric intake itself can significantly affect glucose metabolism[23–25], which could be a potential confounding factor in interpreting metabolic outcomes in pre-clinical cancer cachexia models and in the clinic.

In this study, we aimed to investigate whole-body and skeletal muscle-specific glucose metabolism across two models of pre-clinical cancer cachexia and how short-term cachexia-mimicking food restriction may be a confounding factor in the interpretation of the metabolic effects of cachexia.

## 2. Methods

### 2.1 Animal experiments

All experiments were approved by the Danish Animal Experimental Inspectorate (License: 2021-15-0201-01104 and 2021-15-0201-01085). Mice were maintained under a 12:12 h light/dark photocycle at ambient (22 ± 1 °C) temperature with nesting material. All mice received a rodent chow diet (Altromin no. 1324; Chr. Pedersen, Denmark) and water *ad libitum*. Male mice were group-housed for cancer studies, and single-housed for the food-restriction study.

### 2.2 Tumor-bearing mouse models

12-week-old male Balbc/J mice (Janvier, Denmark) were used for Colon26-cancer (C26) studies, and 14-week-old male C57BL/6JBomTac (Taconic, Denmark) were used for KPC studies. All mice were group-housed in pairs and randomly assigned to either the cancer or control group. C26 adenocarcinoma cells (kind gift from Dr. Adam Rose) were maintained at 37°C, 5% CO2, in RPMI 1640 medium (Gibco, #11875093) supplemented with 10 % fetal bovine serum (FBS, Gibco #F0804) and 1% antibiotic-antimycotic (ThermoFisher, #15140122). KPC cells (CancerTools, #153474) were maintained at 37°C, 5% CO_2_, in DMEM (Gibco, #11965092) supplemented with 10 % FBS (Gibco #F0804) and 1% antibiotic-antimycotic (ThermoFisher, #15140122). All mice were shaved on the right flank the day before cancer inoculation. On the day of inoculation, C26 or KPC cells were trypsinized and washed twice with PBS (centrifuged for 3 min at 1,600 g), followed by resuspension in PBS. Next, all mice were injected subcutaneously into the flank with PBS with or without C26 or KPC cells for a final concentration of 5×10^5^ cells or 1×10^6^, respectively. Mice were terminated when reaching humane endpoints following the tumor guidelines from the Danish Animal Experimental Inspectorate (tumors >14 mm - average length and width or ulcerations >5 mm). C26-cancer mice were divided into non-cachexia (tumor with no weight loss) or cachexia (tumor with weight loss) groups.

### 2.3 Food restriction, pair-fed to cachectic C26 tumor-bearing mice

12-week-old male Balbc/J mice (Janvier, Denmark) were used for the pair-feeding study. All mice were single-housed, and food intake was recorded daily during the last week of the experiment. The last three days, the food intake was reduced by approximately 30%, corresponding to what was observed in cachectic C26-cancer mice.

### 2.4 Body composition

Body mass composition was assessed by nuclear magnetic resonance using an EchoMRI™ (USA) or a Bruker LF90II Body Composition Analyser (Bruker) for total body, lean, and fat mass. All mice were MRI scanned before cancer cell inoculation and the day before termination. The tumor mass from dissections was subtracted from the total body weight and lean mass.

### 2.5 Grip strength

Grip strength was assessed in the week before termination. Grip strength assessment was performed using a grip meter (Bioseb). The mice were held by the base of the tail and briefly left to hover over the grid and allowed to grab with all four paws. Once grabbed, the mice were gently pulled back in a horizontal position. This was repeated three times for each mouse, and the maximal recording was used as final force readout.

### 2.6 Glucose tolerance test

All mice were fasted for 4 hours before the glucose tolerance test (GTT). D-mono-glucose (2g kg^−1^ bw) was administered via intraperitoneal injections, and blood was collected from the tail vein. Blood glucose levels were determined at timepoints 0, 20, 40, 60, 90, and 120 minutes using a glucometer (Bayer Contour, Bayer, Switzerland). Blood aliquots from timepoints 0 and 20 min were centrifuged at 13,000 rpm for 5 minutes at 4°C, and plasma was collected and frozen in liquid nitrogen. Insulin levels were measured using the Mouse Ultrasensitive Insulin ELISA (#80-INSMSU-E01ALPCO Diagnostics, USA).

### *2*.*7 Ex vivo* insulin-stimulated glucose uptake

Soleus and extensor digitorum longus (EDL) muscles were rapidly isolated from anesthetized mice, and non-absorbable 4–0 silk suture loops (Look SP116, Surgical Specialties Corporation) were attached at both ends as previously described[12]. In brief, the muscles were placed in a DMT Myograph system (820MS; Danish Myo Technology, Hinnerup, Denmark) and placed under resting tension (4-5 mN at 30°C) and oxygenated (95% O_2_, 5% CO_2_) Krebs-Ringer buffer. Isolated muscles were pre-incubated in Krebs-Ringer buffer for 5 minutes, followed by 20 minutes of maximal insulin stimulation (60 nM). During the last 10 minutes, 0.75 µCi/mL [3H]-2DG and 0.225 µCi/mL [14C]-Mannitol were added (including insulin). Following the insulin stimulation, muscles were immediately rinsed in ice-cold saline and snap-frozen. Muscle-specific 2DG-uptake was measured on tissue lysates[26].

### 2.8 Lysate preparation and immunoblotting

Muscle tissues were homogenized in a modified GSK3-buffer and prepared for immunoblotting as previously described in detail[17].

### 2.9 Statistical analysis

Data are presented as mean ± SEM and individual data points (when applicable) and analyzed using GraphPad Prism 10. Statistical tests were performed using paired/non‐paired *t*‐tests or repeated/no‐repeated two‐way analysis of variance (ANOVA) as applicable. Multiple‐repeated two‐way ANOVAs were performed in analyses, including all experimental groups testing for the effect of C26, KPC, or food restriction. Sidak’s post hoc test was performed when ANOVA revealed significant main effects and interactions. The significance level was set at α = 0.05.

## 3. Results

### 3.1 C26-cachectic mice exhibit improved glucose tolerance and elevated skeletal muscle insulin-responsiveness *ex vivo*

We first determined classical manifestations of cachexia in the C26-cancer model to establish the effects of cachexia on glucose metabolism. Mice were divided into a control group (n=12), a C26 non-cachectic group (n=6, <10% weight loss), and a C26 cachectic group (n=9, >10% weight loss) based on the weight loss at termination (Fig. 1A). The cachectic mice had a 20% reduction in body mass 22 days after inoculation, while non-cachectic mice maintained their body mass. Control mice had a minor body mass gain of 5% (Fig. 1B). Tumor size was approximately 80% larger in cachectic mice compared to non-cachectic mice (Fig. 1B). Similarly, cachectic mice displayed a 15% reduction in lean body mass, whereas both non-cachectic and control mice maintained lean body mass (Fig. 1C). Cachectic mice exhibited a 75% reduction in fat mass, while non-cachectic (p=0.0510) and control mice exhibited a ∼20% increase in fat mass (Fig. 1D).

**Figure 1:**
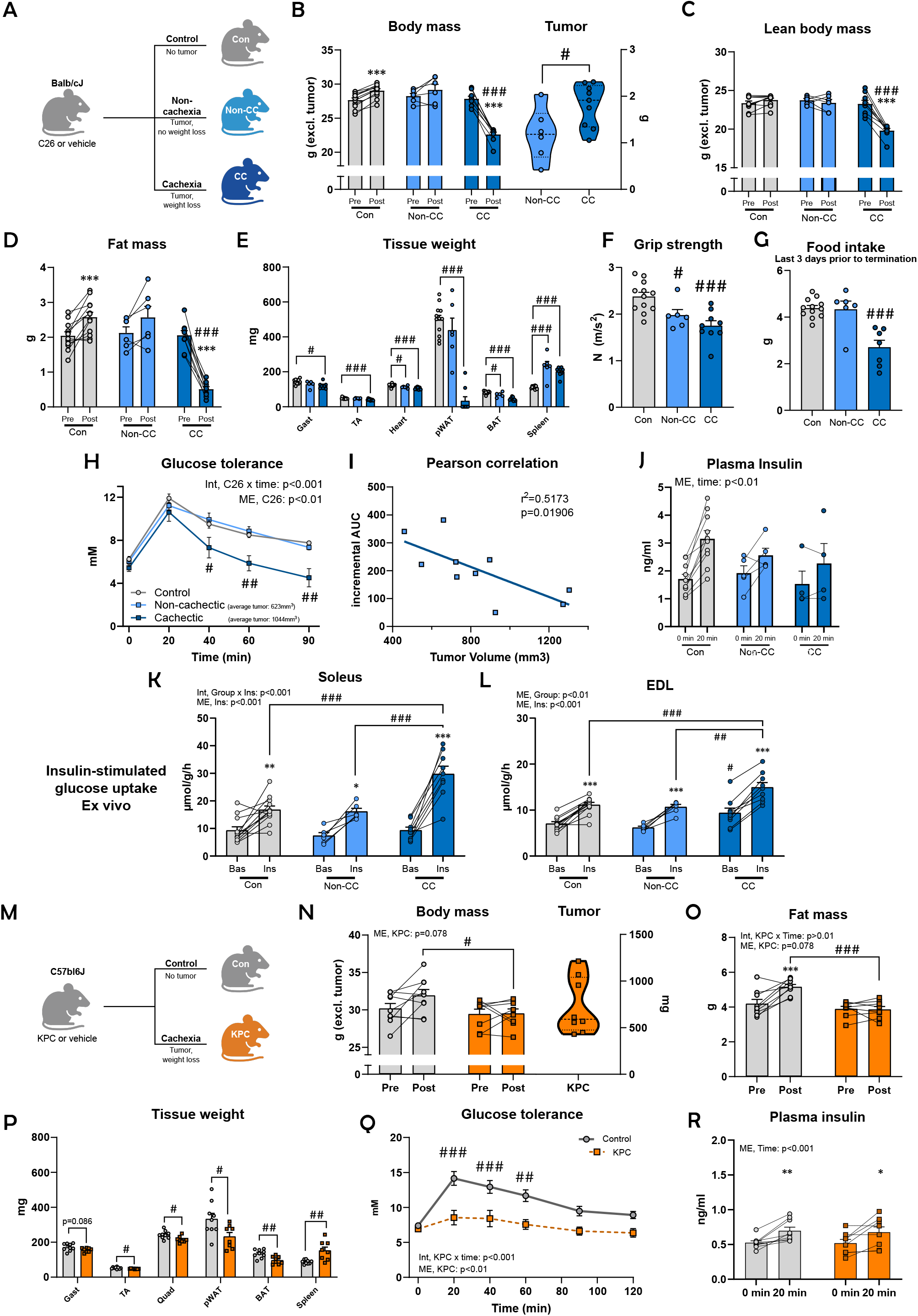
C26 and KPC tumor-bearing mice exhibit improved glucose tolerance despite lowered muscle and fat mass. **A)** Study design of the C26-cancer study. Mice were divided into three groups. 1) Control (n=12), 2) pre-cachectic (tumor, no weight loss, n=6), and 3) cachectic (tumor, weight loss, n=9). **B)** Body mass and tumor mass. **C)** Magnetic Resonance Imaging-derived lean body mass. **D)** Magnetic Resonance Imaging-derived fat mass. **E)** Tissue weights at termination from gastrocnemius (Gast), tibialis anterior (TA), heart, perigonadal white adipose tissue (pWAT), brown adipose tissue (BAT), and spleen. **F)** Grip strength measured on day 19-22 before termination. **G)** Food intake the last three days before termination. **H)** Glucose tolerance test on day 14 after inoculation. C26-tumor-bearing mice were divided into non-cachectic small tumors (<700 mm^3^) and cachectic large tumors (>700 mm^3^). **I)** Pearson correlation analysis of tumor size and incremental area under the curve. **J)** Plasma insulin levels from time-point 0 and 20 min during the glucose tolerance test. Of not, the sample number is lower due to lack of plasma collected during the tolerance test. **K)** *Ex vivo* insulin-stimulated (60 nM) glucose uptake in the soleus muscle. **L)** *Ex vivo* insulin-stimulated (60 nM) glucose uptake in the EDL muscle. **M)** Study design of the KPC-cancer study. **N)** Body mass and tumor mass. **O)** Magnetic Resonance Imaging-derived fat mass. **P)** Tissue weights at termination from Gast, TA, quadriceps (Quad), pWAT, BAT, and spleen. **Q)** Glucose tolerance test on day 30 after inoculation. **R)** Plasma insulin levels from time-point 0 and 20 min during the glucose tolerance test. Abbreviation: C26 = Colon26, Con = control, CC = cancer cachexia, Int = interaction, ME = main effect, AUC = area under the curve. Values are shown as mean ± SEM, including individual values, and as mean ± SD when individual values are not shown. Effect of C26 or KPC: *= p<0.05, **=p<0.01, ***=p<0.001.

At termination, we weighed selected muscles and adipose tissues. Only cachectic mice had lower weights for gastrocnemius (Gast, -17%), tibialis anterior (T. anterior, -18%), and perigonadal adipose tissue (pWAT, -95%) compared to control mice. Heart tissue weights were reduced in both tumor-bearing groups by 8% and 14% for non-cachectic and cachectic mice, respectively. Brown adipose tissue (BAT) was 17% and 47% lower for non-cachectic and cachectic mice, respectively, compared to control mice. The spleen was similarly increased in both tumor-bearing groups by approximately 100% (Fig. 1E), indicative of similar systemic inflammatory burden despite marked differences in tumor mass and cachectic phenotype. Despite no significant reduction in muscle weights, we detected a 17% reduction in grip strength in non-cachectic mice, while cachectic mice showed a reduction of 27% compared to control mice (Fig. 1F). Food intake was assessed three days prior to termination and a reduction (∼40% vs. controls) was observed exclusively in cachectic mice (Fig. 1G).

Considering the reports of glucose intolerance in mice with cancer[15], we performed a glucose tolerance test 14 days after cancer inoculation to assess the effects of cachexia. Dividing C26 tumor-bearing mice into non-cachectic mice (tumor volume <700 mm^3^) and cachectic mice (tumor volume >700 mm^3^), we were surprised to observe increased glucose tolerance in the cachectic mice compared to control and non-cachectic mice (Fig. 1H), contrasting previous findings[11,27]. Glucose excursions during the GTT were inversely correlated with tumor volume, potentially reflecting cachexia-associated metabolic remodeling or tumor glucose sequestration (Fig. 1I). The observed changes in glucose tolerance were independent of glucose-stimulated plasma insulin (Fig. 1J).

Since skeletal muscle is the primary site for glucose disposal, we proceeded to determine insulin-stimulated glucose uptake in soleus and EDL *ex vivo*. In the soleus muscles, insulin led to increased glucose uptake in all groups (Fig. 1K). However, cachectic mice showed ∼80% higher response to insulin non-cachectic and control mice (Fig. 1K). Similarly, in the EDL muscle, cachectic mice shoed an 35-40% elevation in insulin response compared to non-cachectic and control mice (Fig. 1L). Thus, cachexia was associated with elevated insulin responsiveness towards glucose uptake in mouse muscle.

Together, our results demonstrate that cachexia in C26 tumor-bearing mice was associated with improved whole-body glucose tolerance and muscle insulin-stimulated glucose uptake concurrently with muscle and fat mass loss.

### 3.3 The cachectic KPC tumor-bearing mice also display markedly elevated glucose tolerance

Having observed elevated glucose tolerance in the C26 cachectic model, we next assessed glucose tolerance in another pre-clinical cachectic model, the KPC model (Fig. 1M). KPC tumor-bearing mice had an 8% lower body mass at termination compared to control mice, indicative of cachexia (Fig. 1N). Similarly, fat mass was 25% lower in cachectic mice at termination compared to control mice (Fig. 1O). Measurements of tissues at termination revealed that cachectic KPC mice had lower muscle and adipose tissue weight compared to control mice (Gast -10% p=0.081, T. anterior -9%, quadriceps -11%, pWAT -30%, BAT -28%), while the spleen weight was increased by 75% in cachectic mice (Fig. 1P). On day 30 after inoculation, we performed a glucose tolerance test, and similar to the observation in C26 tumor-bearing mice, and in line with previous literature of the KPC model[28], KPC tumor-bearing mice had a marked improvement in glucose tolerance (Fig. 1Q). The change in glucose tolerance was independent of glucose-stimulated plasma insulin (Fig. 1R). Thus, two different pre-clinical cachectic mouse models exhibited elevated glucose tolerance, which could be, at least in part, driven by elevated insulin-responsiveness in skeletal muscle of cachectic mice.

### 3.4 Elevated Akt signaling coincides with higher insulin-stimulated glucose uptake in C26 cachectic mice

After demonstrating an evidently higher muscle insulin responsiveness, we proceeded to investigate intramyocellular insulin signaling (Fig. 2A). We found a reduction in protein contents of Glucose transporter 4 (GLUT4) (-20%), Glycogen Synthase (-13%, p=0.057), Akt2 (-22%, p=0.057), p70S6K (-17%), and rS6 (-29%, p=0.068) in C26 cachectic compared to control mouse soleus muscle. In contrast, Hexokinase II, Pyruvate Dehydrogenase, TBC1D4, GSK3β, and PRAS40 were unaffected (Fig. 2B). It is well known that insulin acts as an anabolic hormone to activate the Akt signaling pathway stimulating glucose uptake and glycogen synthesis in skeletal muscle[29]. Thus, we investigated Akt phosphorylation and downstream targets in response to insulin. In control and C26 non-cachectic mice, we observed a 2-fold and 2.5-fold increase in phosphorylated (p) AKT^thr308^, respectively, in the insulin-stimulated soleus muscle. Yet, in cachectic mice, there was an even higher 4.4-fold increase by insulin compared to the non-stimulated muscle (Fig. 2C). Correspondingly, pAKT^ser473^ was increased by 3.8-fold upon insulin stimulation in control and non-cachectic mice, while cachectic mice exhibited a remarkable 8.5-fold increase (Fig. 2D). Yet we observed a similar increase (3.3- and 3.9-fold) in pTBC1D4^thr642^, a downstream target of AKT, in insulin-stimulated soleus muscle from non-cachectic and cachectic mice, respectively, compared to a 2.4-fold increase in control mice (Fig. 2E).

**Figure 2:**
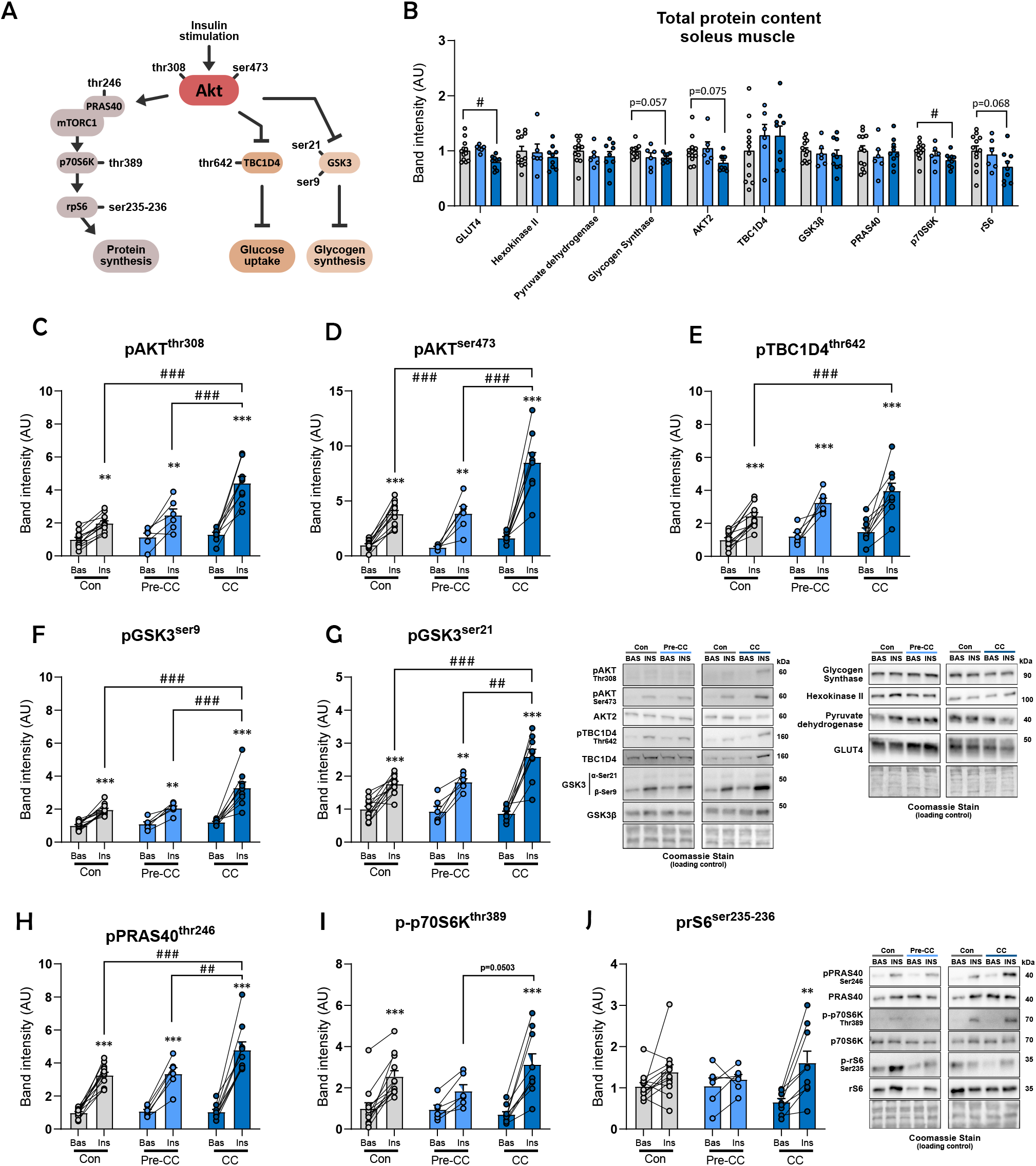
Akt signaling likely drives improvements in *ex vivo* insulin-stimulated glucose transport. **A)** Insulin-stimulated Akt signaling in skeletal muscle. **B)** Total protein content in the soleus muscle of GLUT4 (Thermo Fisher Scientific, #PA1-1065), Hexokinase II (Cell Signaling Technology, #2867), Pyruvate dehydrogenase (D.G. Hardie (University of Dundee, Scotland)), Glycogen synthase (Gift from Prof. Oluf Pedersen), Akt (Cell Signaling Technology, #3063), TBC1D4 (Abcam, ab189890), GSK3β (BD Bioscience, #610202), PRAS40 (Cell Signaling Technology, #2691), p70S6K (Cell Signaling Technology, #2708), rS6 (Cell Signaling Technology, #2217). Protein content is displayed in arbitrary units (AU). Immunoblotting of phosphorylated (p) proteins in the soleus muscle of: **C)** pAKT^Thr308^ (CST, cat#9271), **D)** pAKT^Ser473^ (CST, cat#9271), **E)** pTBC1D4^thr642^ (Cell Signaling Technology, #8881), **F)** pGSK3^ser9^ (Cell Signaling Technology, #9331), **G)** pGSK3^ser21^ (Cell Signaling Technology, #9331), **H)** pPRAS40^thr246^ (Cell Signaling Technology, #2997), **I)** p-p70S6K^thr389^ (Cell Signaling Technology, #9205), **J)** prS6^ser235-236^ (Cell Signaling Technology, #2211). Representative blots are represented with Coomassie staining as a loading control. Values are shown as mean ± SEM, including individual values. Effect of C26: *= p<0.05, **=p<0.01, ***=p<0.001.

Thus, the elevated insulin-responsiveness in cachectic muscle exists despite a reduction of the main glucose transporter (GLUT4), likely circumvented by elevated AKT signaling.

### 3.5 Glycogen and mTORC1 signaling are retained in insulin-stimulated soleus muscle from C26 cachectic mice

Akt phosphorylates several downstream targets. In the insulin-stimulated soleus muscle from cachectic mice, we found a higher increase in pGSK^ser9^ (3.3-fold, Fig. 2F) and pGSK^ser21^ (2.6- fold, Fig. 2G) compared to control and non-cachectic mice (2.0- and 1.8-fold, respectively). PRAS40 is a direct downstream target of AKT during insulin stimulation[30,31]. We detected an elevated abundance of pPRAS40^thr246^ in insulin-stimulated cachectic soleus muscles (4.8- fold), compared to both control and non-cachectic mice (3.3-fold) (Fig. 2H). Insulin-induced activation of the mTORC1 signaling pathway has been widely implicated in the promotion of muscle growth. Despite the presence of cachexia, mTORC1 signaling remained intact, shown by increased p-p70S6K^thr389^ (3.3-fold) and prS6^ser235-236^ (1.6-fold), which contrast previous results in the cachectic *Apc*^Min/+^ mouse model showing reduced mTORC1 signaling after a glucose challenge[32]. Control mice exhibited increased p-p70S6K^thr389^ (2.5-fold), but not prS6^ser235-236^ (Fig. 2I and J). The parallel activation of PRAS40, p70S6K, and rS6 suggests that anabolic signaling downstream of mTOR remained partially intact despite muscle wasting, indicating that cachectic muscle retains the capacity for insulin-induced anabolic responses under these conditions.

### 3.6 Three days of food restriction lowers body weight, muscle mass, and fat mass in mice

We next sought to delineate the underlying contributors to enhanced insulin responsiveness. Anorexia is a key contributor to the manifestations of cachexia, and lowered food intake is known to impact metabolism[23–25]. We therefore executed a 3-day food restriction study (- 30%, mimicking the reduced food intake in C26 cachetic mice) in male BALB/cJ mice to assess whether reduced food intake could explain the improved insulin responsiveness observed in cachectic mice (Fig. 3A). First, we measured body weight daily and discovered that food-restricted mice lost 1.2g (-3,9%) body weight on average after 3 days, compared to control mice who gained 0.7g (2.1%) body weight (Fig. 3B). We measured tissue weights and observed a reduction in tissue weights of all skeletal muscles and adipose tissues upon food restriction (TA -4%, Gast -7%, Quad -7%, iWAT -34%, pWAT -22%, BAT -23%, spleen -16%), while the heart weight was similar (Fig. 3C). Despite lower muscle weights in food-restricted mice, grip strength was unaffected (Fig. 3D), indicating that anorexia may underlie some body weight loss but does not compromise muscle function in tumor-bearing mice.

**Figure 3:**
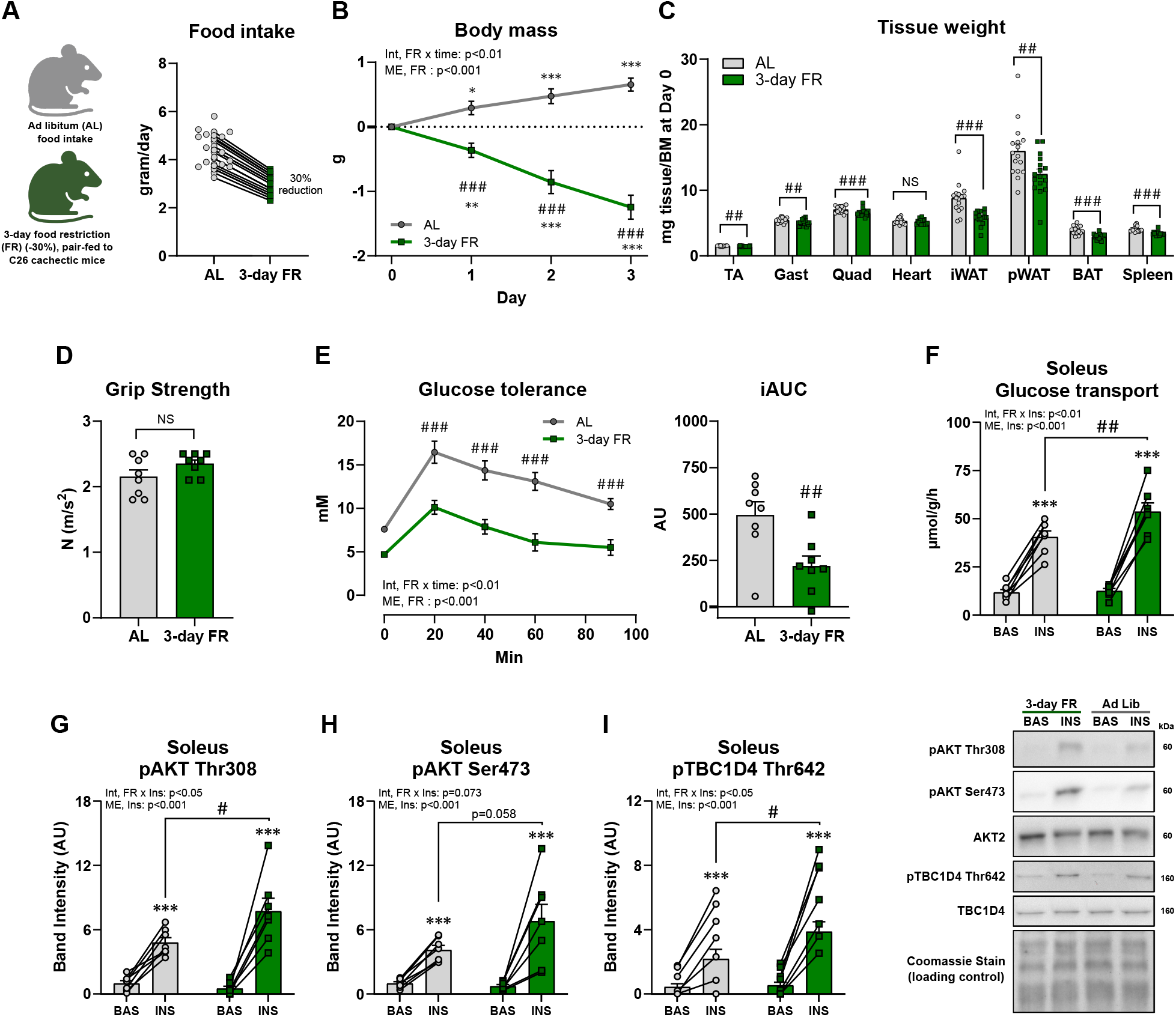
Three days of food restriction in mice improves glucose tolerance, *ex vivo* insulin-stimulated glucose transport in soleus, and enhances Akt signaling. **A)** Study design of the food restriction study. **B)** Body weight measured during the food restriction period. **C)** Tissue weights at termination from TA, Gast, Quad, heart, inguinal white adipose tissue (iWAT), pWAT, BAT, and spleen. **D)** Grip strength measured after 3 days of food restriction. **G)** Glucose tolerance test and incremental area under the curve (iAUC) after three days of food restriction. **F)** *Ex vivo* insulin-stimulated (60 nM) glucose uptake in the soleus muscle. Immunoblotting of phosphorylated proteins in the soleus muscle of: **C)** pAKT^Thr308^ (Cell Signaling Technology, cat#9271), **D)** pAKT^Ser473^ (Cell Signaling Technology, cat#9271), **E)** pTBC1D4^thr642^ (Cell Signaling Technology, #8881). Values are shown as mean ± SEM, including individual values, and as mean ± SD when individual values are not shown. Effect of 3-days food restriction: *= p<0.05, **=p<0.01, ***=p<0.001.

### 3.7 Glucose tolerance, muscle *ex vivo* insulin-stimulated glucose uptake and Akt signaling in increased upon food restriction

To investigate if anorexia may underlie some of the improvements in glucose tolerance and muscle insulin responsiveness, observed in cachetic C26 mice, we determined glucose tolerance and muscle insulin reonsiveness in 3-day food-restricted Balb/c mice. Interestingly, food-restriction markedly improved glucose tolerance with a 56% reduction in iAUC compared to ad libitum fed control mice (Fig. 3D). We then proceeded to determine *ex vivo* insulin-stimulated glucose uptake in the soleus muscles. We were intrigued to see that food-restricted mice had a higher insulin response (30%) compared to control mice (Fig. 3F). In line with our observations in C26 cachectic mice, AKT signaling was elevated in food-restricted mice. This was evidenced by, pAKT^thr308^ increasing 8-fold in response to insulin in food-restricted mice compared to 5-fold in control mice (Fig. 3G). Likewise, pAKT^ser473^ increased by 7-fold in response to insulin in food-restricted mice compared to 4-fold in control mice (Fig. 3H). In addition, we found a 6-fold increase by insulin in pTBC1D4^thr642^ in food-restricted mice, which was only increased by 3-fold in control mice (Fig. 3I).

Taken together, our findings show that reduced food intake is a potential driver of enhanced glucose tolerance and skeletal muscle insulin responsiveness in cachectic C26-cancer mice, associated with enhanced Akt signaling.

## 4. Discussion

In this study, we investigated glucose metabolism and skeletal muscle insulin responsiveness, using two distinct tumor models, C26 and KPC, and in pair-fed food-restricted mice. Our results highlight three key discoveries. First, cachexia was associated with increased glucose tolerance in both C26- and KPC tumor-bearing mice. Second, cachectic, but not non-cachectic, C26 mice exhibit elevated skeletal muscle insulin responsiveness associated with increased Akt signaling. Third, food restriction, mimicking the lower food intake in C26 cachectic mice, caused a metabolic phenotype phenocopying cachexia. These findings suggest that decreased food intake contributes to elevated insulin-responsiveness in cancer cachexia, illuminating an intriguing interplay between food intake and cachexia-associated metabolic reprogramming.

Our first key finding demonstrates that cachectic C26-tumor-bearing mice exhibited improved glucose tolerance compared to non-cachectic mice, which was corroborated in cachectic KPC-tumor-bearing mice. Here, we identified a relationship between tumor burden and glucose tolerance, with cachectic mice harboring significantly larger tumors. It is plausible that larger tumors dispose more glucose during hyperglycemia, contributing to improvements in glucose tolerance. A marked metabolic change in glucose utilization is supported by a recent study, using isotopic tracers in C26 cachectic mice, showing that one-carbon metabolism was a tissue- overarching pathway characterized in wasting leading to glucose hypermetabolism[19]. However, our second key finding highlights that cancer-associated cachexia is also accompanied by elevated skeletal muscle insulin responsiveness. Thus, these data suggest that changes within the skeletal muscle likely contribute to the enhanced glucose tolerance in cachectic mice.

In the soleus and EDL muscles, insulin responsiveness was markedly greater in cachectic compared to both control and non-cachectic tumor-bearing mice. This was observed despite a reduction in protein content of the key glucose transporter during insulin stimulation, GLUT4[33]. Yet, increased GLUT4 translocation to the muscle membrane was indicated by a substantially elevation of phosphorylation AKT (Thr308 and Ser473) in response to insulin. Thus, the current data support the idea of cancer cachexia being associated with elevated glucose tolerance and insulin responsiveness, although the molecular mechanisms remain to be defined.

Our last key finding was that reduced food intake, mimicking food intake in cachexia, could contribute to the increased insulin responsiveness in our cachectic mice as a compensatory adaptation to nutrient limitation arising from anorexia and tissue loss. In our C26 model, only cachectic mice had significantly reduced food intake (∼40 %) with severe lean and fat mass loss. In this context, improved insulin-stimulated glucose uptake in muscle may reflect heightened responsiveness to insulin when exogenous glucose availability is limited, a phenomenon reminiscent of the known effects of caloric restriction[34–36]. Notably, while improved insulin sensitivity may initially represent a compensatory mechanism to preserve energy efficiency during anorexia, chronic activation of this pathway in the face of catabolic conditions such as cachexia fails to prevent muscle loss. Underscoring this is our finding that cachexia-mimicking food restriction alone did not alter muscle strength measured by grip strength in the current study. Decreased grip strength was only observed in tumor-bearing mice and in both cachectic and non-cachectic mice, in line with another study showing decreased muscle function in C26 before cachexia [37]. Accordingly, it is also known that decreased energy intake in cancer is not the sole driver of cachexia [38,39]. Thus, while decreased food intake likely affects the systemic metabolism in cancer cachexia, the pathology of cachexia is complex and involves multiple factors[38].

## 5. Conclusion

In summary, our study reveals that improved glucose tolerance and enhanced skeletal muscle insulin responsiveness in cachectic C26-cancer mice are likely affected by anorexia-induced adaptations. Our data highlight enhanced Akt signaling as a key node linking nutrient status to skeletal muscle glucose metabolism during cancer progression. These findings challenge the assumption that cachexia universally associates with insulin resistance and underscore the need to consider both cancer type, cancer progression, and food intake when interpreting metabolic outcomes in cancer.

## Acknowledgement

We thank our colleagues at the Molecular Metabolism in Cancer and Aging Group, Faculty of Health and Medical Sciences, University of Copenhagen, for profitable discussions on this topic. We acknowledge the Rodent Metabolic Phenotyping Platform (RMPP), Novo Nordisk Foundation Center for Metabolic Research, University of Copenhagen, for the use of their facilities. We also acknowledge Simon Bech Petersen, Department of Biomedical Sciences, Faculty of Health and Medical Sciences, University of Copenhagen, for his assistance. Illustrations were generated using Biorender.com (RRID:SCR_018361) and ©Inkscape. Graphs were generated with GraphPad PRISM (RRID:SCR_002798).

## CRediT

Conceptualization: EF, LS, SHR

Data curation: EF, KWP, ZKJO, PP, JRK, LS, SHR

Formal analysis: EF, SHR

Funding acquisition: LS, SHR

Investigation: EF, KWP, ZKJO, PP, JRK, LS, SHR

Methodology: LS, SHR

Project administration: EF, LS,

SHR Resources: LS, SHR

Supervision: LS, SHR

Validation: EF, SHR

Visualization: EF, LS, SHR

Writing – original draft: EF, LS, SHR

Writing – review and editing: all authors

## Declaration of generative AI and AI-assisted technologies in the writing process

Statement: During the preparation of this work, the authors used ChatGPT to do minor corrections and shorten sentences. After using this tool/service, the authors reviewed and edited the content as needed and we take full responsibility for the content of the published article.

